# Self-guided search in immersive VR reveals active hierarchical planning

**DOI:** 10.64898/2026.09.23.753863

**Authors:** Qianqian Wan, Shiyang Ren, Shea E. Duarte, Jonathan Park, Charan Ranganath, Jack J. Lin, Simona Ghetti, Joy J. Geng

**Affiliations:** Department of Psychology, University of California, Davis; Center for Mind and Brain, University of California, Davis; Center for Neuroscience, University of California, Davis; Department of Neurology, University of California, Davis

**Keywords:** naturalistic visual search, virtual reality, eye movements, hierarchical planning, active sensing

## Abstract

Despite the ubiquitous nature of visual search in our daily lives, our knowledge of the underlying mechanisms is almost entirely derived from studies using simple, synthetic stimuli briefly flashed on a computer monitor. In real life, however, visual search occurs in three-dimensional environments over much longer timescales and involves the dynamic coordination of attention, memory, and action planning, generating novel predictions that cannot be tested with traditional approaches. Here, we examined how self-guided visual search unfolds under naturalistic conditions that extend over space and time. We asked participants (*N*=80) to find specific objects (e.g., a green lamp, an orange shelf) in an immersive, visually rich virtual reality environment using range-limited teleportation (2 m steps). Consistent with our hypotheses, we found that self-guided search engaged a hierarchical information-seeking policy characterized at each stage by recurring plan-and-execute cycles paired with subgoal-specific attentional priorities. Subgoal priorities progressed from wayfinding to identification of clusters of target-similar objects, and then to specific target features. These search priorities progressively narrowed the search space through coordinated eye, head, and body movements that recurred throughout the task. We conclude that naturalistic search is hierarchical, built from sequential subgoals with shifting attentional priorities that single-level tasks cannot capture. Moreover, immersive VR with eye-head-body tracking dissociates motor planning from execution moments and reveals how looking content shifts with each task subgoal while the person moves through space, something desktop tasks cannot capture.

**Significance Statement:** Most of what we know about how people find things comes from simple images on computer screens, yet everyday search unfolds as we move through cluttered, three-dimensional spaces. Therefore, our models of visual search need to account for this complexity. Using immersive virtual reality that tracked the eyes, head, and body, we show that natural search is organized as a hierarchy of subgoals rather than one sweep for the target. People repeatedly paused to plan and then acted, and what they sought shifted in order, first finding a route, then clusters of promising objects, and finally specific target features. This structured strategy is invisible to flat laboratory tasks, and it opens a way to characterize attention, memory, and action as they interact in the real world.

## Introduction

Imagine going to buy a green desk lamp at a large department store. From the store entrance, you must first find the lighting fixture area, then locate the section with desk lamps, before finally picking out the specific lamp you want. To understand how we find objects, typical experimental visual search tasks use 2D desktop computers with simple, briefly presented, synthetic stimuli that require only a few saccadic eye movements before target verification^1–6^.

However, naturalistic search likely requires multi-step goals and action planning over the course of minutes to hours. This apparent complexity raises questions about whether simple desktop tasks provide a sufficient account of how self-guided search occurs in daily life.

One possibility is that naturalistic search can be understood based on the sequence of steps required to complete simpler search paradigms, only requiring holding target information continuously in memory for longer. From this perspective, current approaches may suffice to characterize visual search in naturalistic environments ^7^. Alternatively, we hypothesize that search in naturalistic environments is better characterized as a hierarchical plan in which a series of subgoals defines the current set of attentional priorities used to identify information just-in-time for the next set of actions (Figure 1). Each subgoal should reflect its unique set of attentional priorities to identify the best next action, which will progressively narrow the search space and bring the observer closer to the final target ^8^. If the latter is true, we should expect eye-gaze to be selectively directed to information relevant to the current subgoal, especially in moments when action planning occurs. Body movements also have to be coordinated with the eyes to find the vantage points that can inform the upcoming decisions. How this occurs in a naturalistic search, however, remains unknown.

**Figure 1.**
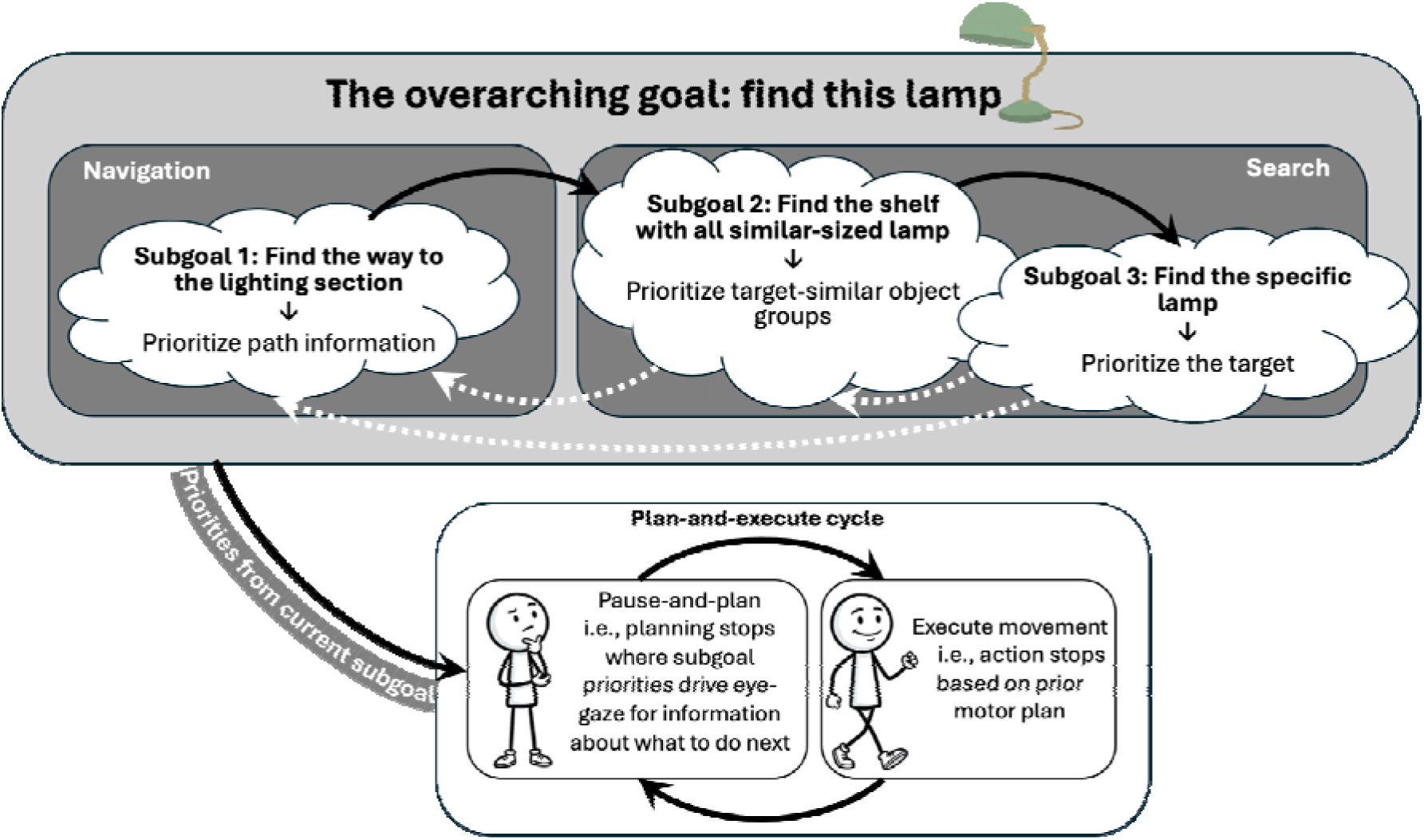
Conceptual model of search task hierarchy. To find the green desk lamp efficiently, one must complete several subgoals one by one, with recurring cycles of planning and movement execution (i.e., the plan-and-execute cycle). Importantly, each subgoal defines a specific set of attentional priorities for the planning moments: *Subgoal 1* involves sampling information about alternative paths to decide how to get to the target display room; *Subgoal 2* prioritizes information for moving closer to likely target locations, without requiring the movement to be definitively towards the target; and *Subgoal 3* focuses on locating the specific target for pick-up. The order in which subgoals are executed depends on the current environment, i.e., failure at *Subgoal 3* might lead one to go back to *Subgoal 2*, but each must be engaged sequentially.

One way that subgoal planning might be instantiated is through physical pauses paired with increased head and eye-gaze movements to allow for information gathering as observed in rodents and humans during navigation tasks^9–20^. The active sensing framework provides an account of this relation between motor actions and information seeking based on internal goals ^21^: Active sensing situates a person within a closed-loop system in which current sensory information guides the creation of action plans designed to acquire new sensory information that further reduces task uncertainty^22–32^. The sensory information gathered by eye movements is specifically tuned to information immediately relevant for the next body movement, rather than more temporally distal ones^19,33–37^. This literature suggests that looking behaviors scaffold the next action by extracting precise visual information to guide motor control, while at the same time, motor actions optimize sensory processing ^8^. However, no study to date has tested this eye-head-body coordination in a naturalistic, self-guided search task where action plans must be constantly updated across different subgoals.

On this basis, we predict that there will be common physical behaviors related to stopping and head swivels associated with cognitive planning that occur throughout our task. However, the exact information sought during these moments with eye-gaze should be specific to the current subgoal within the overarching task goal. To return to our opening example, when navigating to the lighting fixture area at the store, we expect that planning moments during the first subgoal (i.e., finding the lamp room) would prioritize attention towards information for wayfinding^38^, whereas planning during subsequent subgoals (i.e., focused on lamp localization and identification) would prioritize any information that brings the actor closer to the final target, such as to an aisle of desk lamps, followed by target-defining features, such as a green desk lamp^7,24,32,39–52^.

To understand how attention operates on the timescale of real-world search, we asked participants to navigate a 4,216 m² furniture store in immersive virtual reality to find a specific target object while recording their eye, head, and body movements. We theorize that search is organized hierarchically, with finding the target as the overarching goal while navigational and search subgoals are pursued in turn within behavioral plan-and-execute cycles (Figure 1). There are two overarching hypotheses guiding this research. First, we hypothesize that plan-and-execute cycles are a common behavioral signature across subgoals, visible as periodic planning moments when participants pause and sample the scene with wider head and eye movements before acting. Second, we predict what attention gathers at those planning moments should be subgoal-specific, shifting from where to go next while navigating to progressively finer target information while searching. Based on classical feature-based guidance models, we would expect that attention would prioritize objects with target-similar color and size equally throughout search, or even favor color as a dominant guiding feature^41,45,46^. In contrast, we hypothesize that attention will prioritize features based on their spatial arrangement first. We make this hypothesis based on the expectation that attentional selection during real-world tasks is bound primarily by the need to spatially triangulate possible target locations and therefore search progresses using coarse-to-fine target-relevant information.

## Results

Eighty participants searched a multi-room virtual furniture store for a set of everyday target objects following an initial exploration period in which they were familiarized with the environment (see Methods). Participants moved through the store by teleporting in range-limited steps of 2 m or less and were free to turn their head and body to look around, while their gaze and head position were recorded (see Methods, Figure 2). Each trial began from the same position at the entrance of the store. Successful target localization required navigating through the store to the target category’s “display room”, where the target could be found.

**Figure 2.**
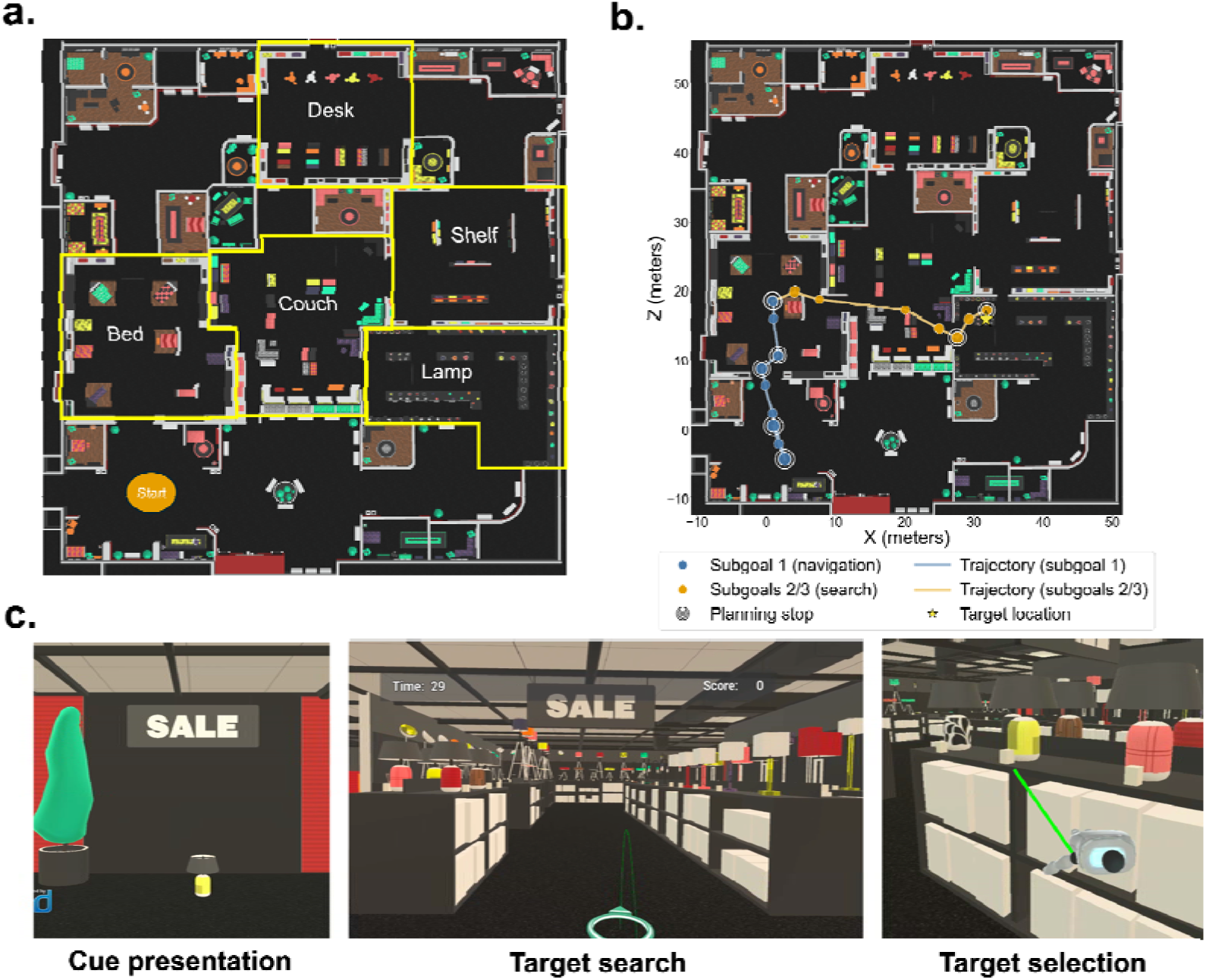
Task design and procedures. **(a)** Bird’s-eye view of the 4,216 m² furniture store, with the five target display rooms labeled. **(b)** An example trial trajectory, with teleportation stops colored by subgoals and outlined by stop type (planning, action), with dot size scaled to stop duration. **(c)** First-person view in VR at different trial stages: cue presentation (6 s), target search and selection (up to 4 minutes). Upon entering the target display room (the middle figure), participants needed to decide whether to approach the more clustered, size-similar objects (as shown in the green circle, not visible to participants) or the more dispersed, color-similar objects (pink circles) and search until the target was found.

We hypothesized that the search task could be decomposed into subgoals related to navigation, target search, and target localization, each with unique attentional priorities.

Moreover, we hypothesized that these subgoal states would be most easily identified during spontaneous moments when people plan their next actions. In what follows, we first describe the plan-and-execute cycle, a putative common signature of planning, and then the unique informational content participants sampled through eye-gaze during each subgoal. We hypothesized that while the body makes similar actions at each planning stop, attention will be predominately guided by the objective of each subgoal.

### The plan-and-execute cycle is a common behavioral signature of planning

Natural search behaviors are expected to include intermittent planning moments that guide the next immediate actions. Here, we used stop duration and head movement with unsupervised clustering algorithms to detect these hypothesized planning stops (when participants are expected to pause to sample the environment and decide where to go next) and distinguish them from action stops (when participants are expected to advance toward a previously chosen region) (see Methods and Supplementary Materials). We successfully identified these two classes of stops based on duration and amount of head movement. To exclude the possibility that these stops reflected behavioral randomness, rather than actual differences in planning, we predicted that high magnitude planning stops would be more likely to cluster at choice points shared across trials, whereas action stops would be more spatially diffuse given that they exploit already-chosen trajectories. By measuring the dispersion of each stop type with the entropy of its binned spatial distribution and comparing it against a permutation null distribution (Methods; Figure 3a), planning stops were found to be significantly more spatially clustered than chance (observed *H* = 10.44, null *M* = 10.59, 95% CI [10.55, 10.63]; *p* < .001), whereas action stops were not (median *H* = 10.63, *p* = .972). This is consistent with the idea that there are common decision points during navigation, likely due to the environment’s configuration, e.g., the location of corners, or junctures with many path options. In addition, we expected that planning stops would be more likely to happen at decision points that lead to changes in behavior. Consistent with this prediction, planning stops tended to be followed by larger changes in heading direction compared to action stops (Figure 3b, see details in Supplementary Materials).

**Figure 3.**
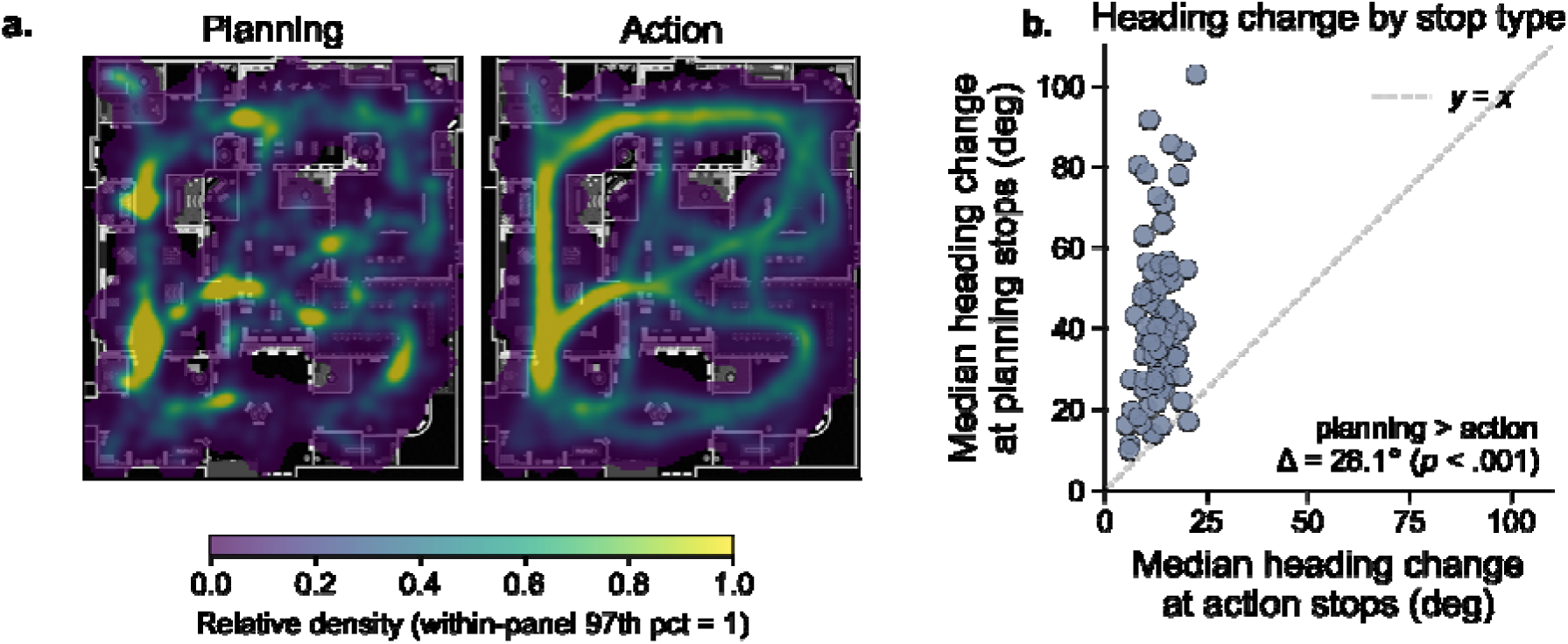
Behavioral signatures of the plan-and-execute cycle. (a) Spatial distributions of planning stops in comparison with action stops. Stop positions were binned into 0.5 m grids, smoothed with a 2D Gaussian, and normalized within the panel. Each cell was scored as visited or not per trial (presence/absence), so repeated stops within one trial could not inflate its density. For visualization, we normalized each panel’s density to its 97th-percentile value (rather than its maximum) and clipped it to [0, 1], a standard robust-scaling step that prevents a small number of high-density cells from saturating the color map; this affects only the color mapping, not any reported statistic. **(b)** Heading direction change comparison. Each point is one participant; the dashed line denotes equality. Points above the diagonal indicate participants whose heading change was larger at planning than at action stops.

Together, we found that stops with longer durations, more head swiveling, and broader looking occurred at specific, choice-relevant junctions and coincided with changes in heading direction. These characteristics are consistent with the profile of a planning stop instantiated similarly across all subgoals.

However, the information used for planning decisions should differ as a function of subgoal: participants primarily need to determine which path to take when navigating (subgoal 1), which object cluster to approach when narrowing the space in the target room (subgoal 2), and finally, identify which object is the actual target for pick-up (subgoal 3). We investigated these attentional priorities during planning using eye-gaze next.

### Subgoal 1 prioritizes alternative path information during planning for navigation

Subgoal 1 is to navigate from the store entry to the target display room. This represents the coarsest subgoal in service of locating the target. In order to accomplish this, individuals must decide which paths to take, i.e., wayfinding, which we hypothesized would prioritize attention to information about path options. Consistent with this hypothesis, we found participants directed more looks towards alternative paths (i.e., navigable routes that were not subsequently taken) than the path-taken (i.e., routes within 10° of the participant’s next movement, see Methods and Figure 4b) during planning stops relative to action stops.

**Figure 4.**
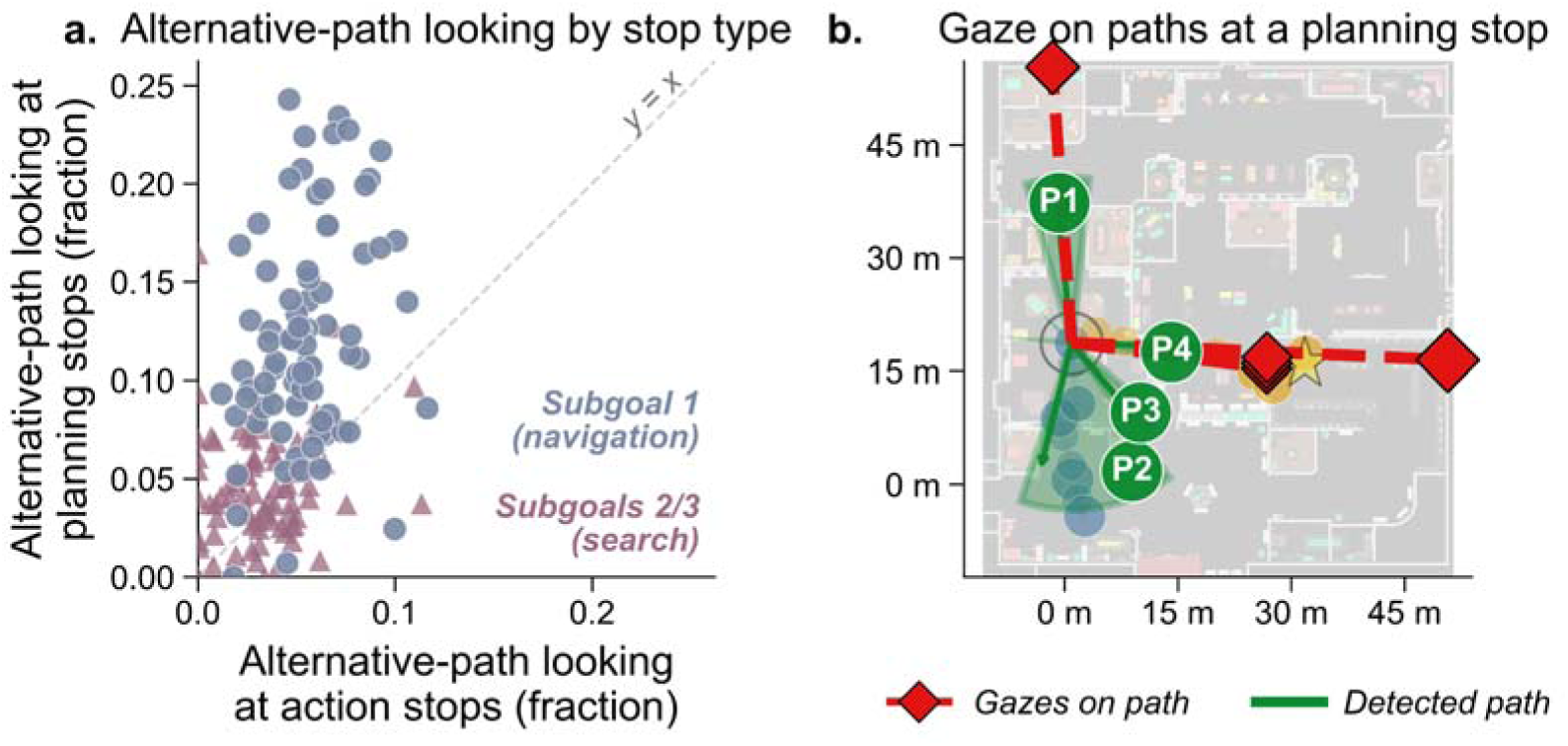
Planning stops preferentially sample alternative paths during initial wayfinding (Subgoal 1). **(a)** Per-subject fixation time on *alternative* paths (paths not subsequently taken), at planning versus action stops, expressed as a fraction of each stop’s total fixation time. Each marker is one participant (*N* = 80); the *y*-axis is the mean fraction at that participant’s planning stops, and the *x*-axis is the mean at their action stops. Dark blue circle markers show planning stops from Subgoal 1 (navigation to the target display room); pale blue triangle markers show Subgoals 2–3 (search inside the room). Points above the dashed *y* = *x* line indicate more alternative-path looking at planning than action stops. **(b)** Illustrative single stop showing an example case of alternative-path looking on the store map at a single planning stop. Green wedges are the alternative paths available from the participant’s viewpoint (P1–P4); red dashed lines with diamonds are gaze episodes directed along a path.

Specifically, we used a linear mixed-effects model in which stop type (planning vs action), path-looking type (path taken vs alternative paths), and their interaction were entered as predictors of arcsine-square-root-transformed path-looking, with target object type as a covariate, with subject-level random intercepts. We found a significant stop type × path looking type interaction, β = 0.20, 95% CI [0.17, 0.23], *p* < .001, such that fixation time on alternative paths was higher at planning stops than action stops (*M* = 0.12 [0.11, 0.13] vs. *M* = 0.06 [0.05, 0.06]; Mann–Whitney U = 5699, Cohen’s *d* = 1.60, *p*_adj_ < .001). In contrast, fixations during action stops concentrated mostly on the path subsequently taken (*M* = 0.27 [0.25, 0.28]; planning vs. action *U* = 1218, Cohen’s *d* = -1.17, *p*_adj_ < .001) (Figure 4a). This shift towards alternative paths at planning stops reflected a redistribution of path looking rather than simply more path looking overall: planning and action stops devoted statistically indistinguishable total time to paths during subgoal 1 (*M* = 0.31 [0.29, 0.33] vs. *M* = 0.32 [0.31, 0.34]; *U* = 2913, Cohen’s *d* = -0.16, *p* = .328). This pattern was specific to the wayfinding subgoal 1; the alternative path looking at planning stops was significantly less during subgoals 2 and 3 (*M_subgoal1_* = 0.12 [0.11, 0.13] vs. Msubgoal2&3 = 0.05 [0.04, 0.05]; Mann–Whitney *U* = 5772, Cohen’s *d* = 1.71, *p* < .001; paired within-subject difference = 0.08 [0.06, 0.09], *p* < .001), suggesting subgoal-specific attentional prioritization.

Although wayfinding was prioritized during navigation, an exploratory analysis showed that the current target’s features also guided gaze before participants reached the display room, indicating that the overall goal was maintained across subgoals rather than entirely set aside within each (see Supplementary Results).

Next, we identified the key transition moment between subgoals 1 and 2 using the first fixation into the target display room. This self-guided moment served as a stable separator between subgoals and emerged from natural behavior. After this first fixation, participants rarely made planning stops until they were inside the display room, with 74.5% of the trials containing no planning stop. On average, the planning stop rate during this transitional period (*M* = 0.13, *SD* = 0.24) was significantly lower than in every other comparison window, *p*s < .001. The lack of planning stops after the first fixation into the target display room and before entering the room suggests this was a consistent boundary moment during task completion. Moreover, once people entered the target display room, they rarely left (92.5% of trials had single, terminal entry), further suggesting that the transition into the target room signaled a new phase of task completion.

### Subgoal 2 prioritizes spatial clustering more than feature target-similarity alone when inside the target display room

We next examined the information sought during planning stops once participants identified the target room. Classical feature-based guidance models would predict that attention should prioritize objects with target-similar color and size^41,45,46^. We sought to verify this and to test our hypothesis on how the spatial arrangement of objects in the room would modulate this effect.

To test this hypothesis, we first verified that the similarity effect was present in our large-scale environment. To do so, we obtained the feature similarity scores for each object in a separate rating task (6 = most similar, 1 = least; see Methods). These ratings were used as a continuous measurement of a single object’s similarity to the target (in size and color). In a binomial generalized linear mixed model predicting the fixation probability of each object, we found feature similarity predicted the fixation likelihood positively (size: *OR* = 1.25, *b* = 0.22, 95% CI [0.20, 0.24], *z* = 23.60, *p* < .001; color: *OR* = 1.16, *b* = 0.15, 95% CI [0.12, 0.18], *z* = 10.63, *p* < .001, see Figure 5a). Having confirmed that target-similar objects are prioritized, we next tested our hypothesis that spatial arrangement would modulate the feature-similarity effect. We call this the clusteredness hypothesis, which stems from our proposal that search proceeds in a hierarchical fashion.

**Figure 5.**
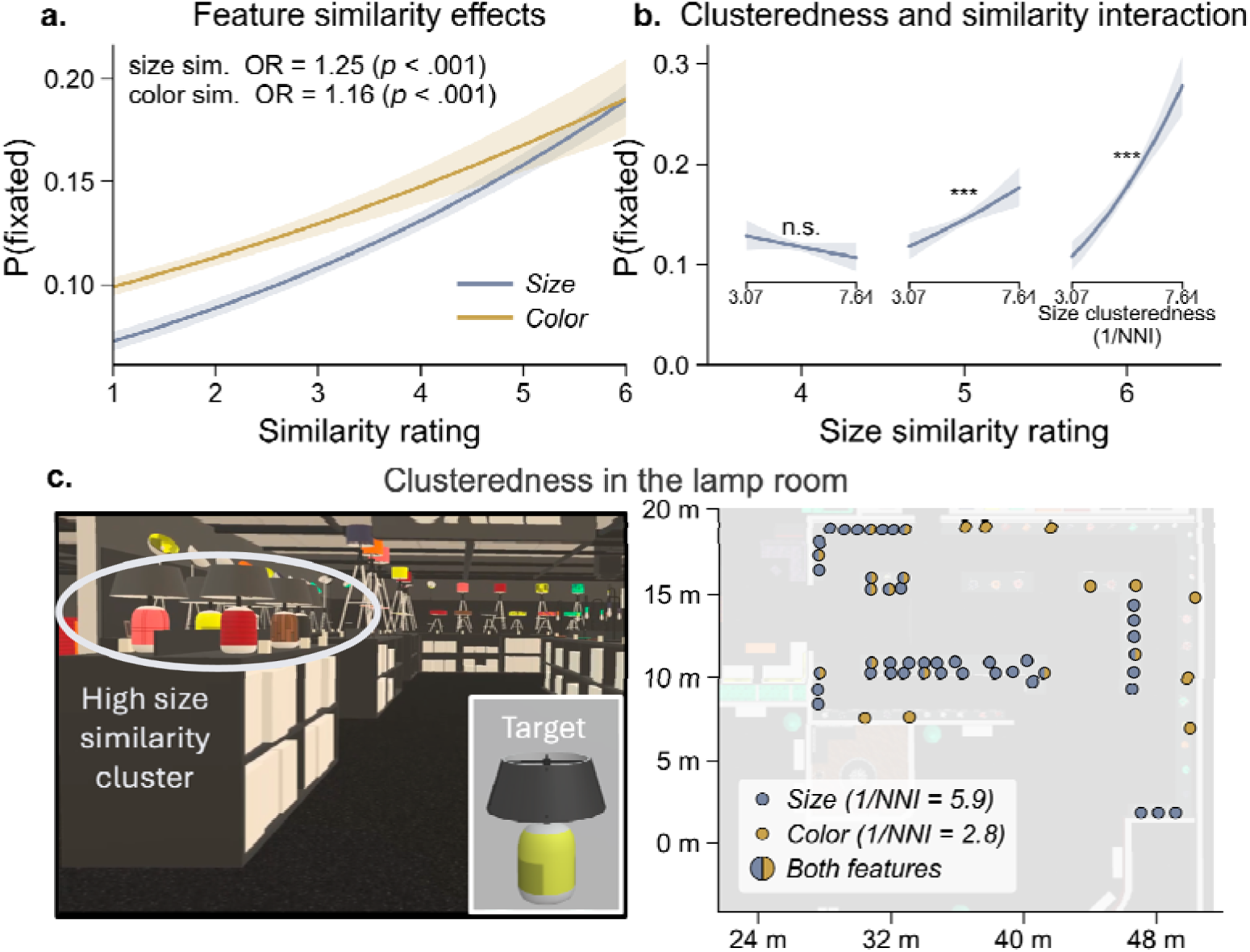
Model-predicted fixation probability and illustration of the clusteredness calculation. **(a)** Feature similarity effects. Predicted fixation probability as a function of its similarity rating, shown separately for size and color similarity, with the other feature held at its mean. Shaded bands are 95% confidence intervals. **(b)** Effect of clusteredness on fixation probability at high similarity ratings. Positive slopes indicate that clustering increases fixation probability. Shaded bands are 95% confidence intervals. **(c)** An example first-person view inside the lamp room (left) and the spatial clusteredness by feature similarity on the lamp room floor plan (right).

Objects in our target display rooms were spatially organized so that similarly sized objects were more likely to cluster together. The spatial cluster provides a center of mass for motor planning for “where to go next.” Approaching a cluster with one target feature, then searching within it for the second feature, is more efficient than approaching physically dispersed single objects one at a time. Therefore, if clusteredness matters for narrowing the search space, we would expect fixation probabilities to increase when objects are in a cluster of other target-similar objects. Alternatively, if attention operates based on feature-based guidance alone, fixation probability should increase with feature similarity and be unaffected by spatial arrangement.

To test this, we created a metric of clusteredness for each object in each target display room. Objects were first divided into two groups based on feature-similarity (high-similarity = rating ≥ 4, or low-similarity = rating ≤ 3). Within each group, we then computed each object’s clusteredness based on its physical proximity to other objects. For example, for an object with a high size-similarity rating, i.e., ≥ 4, clusteredness was calculated using the number and distance of its high-size-similarity neighbors (1/NNI; higher values mean the object is located within a tighter cluster; see Methods and Figure 5c). We then fit a binomial generalized linear mixed model predicting each object’s fixation probability at each planning stop from its size, color, size clusteredness, and color clusteredness, modeling the size and color terms and their interactions separately.

As predicted, a significant interaction between feature similarity and clusteredness was found (size: OR = 1.16, *b* = 0.15, 95% CI [0.12, 0.18], *z* = 10.84, *p* < .001; color: OR = 1.54, *b* = 0.43, 95% CI [0.08, 0.78], *z* = 2.40, *p* = .016; OR stands for odds ratios (*e*), where OR > 1 indicates increased and OR < 1 decreased odds of fixation per one-unit change in the predictor value). As shown in Figure 5b, the probability of fixating an object increased with clusteredness for objects with a size-similarity rating of 5 or 6 (rating 5: OR = 1.11, *b* = 0.10, 95% CI [0.05, 0.16], *z* = 3.79, *p* < .001; rating 6: OR = 1.29, *b* = 0.25, 95% CI [0.19, 0.31], *z* = 8.18, *p* < .001).

This result shows that clusteredness increased the likelihood of looking at an object with high target similarity, consistent with our hypothesis that spatial clusteredness is used to guide search, over and above that of feature-similarity alone. Clusteredness did not increase the probability of fixation for objects with a size-similarity rating of 4 (*b* = −0.05, 95% CI [−0.11, 0.01], OR = 0.96, *z* = −1.51, *p* = .130), reinforcing the notion that clusteredness increased attentional priority only for objects with high target-feature similarity. Since the color feature had overall low clusteredness, the results were not meaningful (but see full model details in the Supplementary Results).

Together, these results suggest that in large search areas, spatially clustered high-similarity features are prioritized during planning stops when decisions are made about where to go next. While the specific effect of size is likely related to the organization of objects in our display rooms, the data highlight the importance of spatial information in constraining naturalistic searches that require movement through dispersed objects.

### Subgoals 2 and 3 unfold temporally inside the target display room

The previous analysis showed that fixations were biased towards the clustered size feature. Next, we wanted to understand how subgoals 2 and 3 temporally unfolded. If spatial clusteredness is related to a search plan that prioritizes broad-to-specific information, we would expect cluster looking (subgoal 2) to occur early in the trial, followed by more specific feature-based looking (subgoal 3) to localize the target (see Supplementary analyses, Figure S8).

Because natural looking is heterogeneous, with fixations at each time point being driven by potentially multiple task-relevant sources of information as well as task-irrelevant looking, we used a hidden Markov model (HMM) to recover the latent states of attentional priority from the local sequence of stops, followed by a regression analysis to identify the sequential probability of each latent state occurring within a trial.

The HMM jointly estimates the number of states needed to describe the sequence (via BIC), each state’s characteristic distribution over looked-at categories (the emission distribution), and the probability of moving from one state to the next (the transition matrix). For this analysis each planning and action stop was summarized by the object category looked at most (target, high target color-and-size similarity, high color similarity, high size similarity, or low similarity), and this per-stop category was the observation the HMM modeled (Methods).

The best-fitting model had four states (BIC = 10,873.37, log-likelihood = −5,307.25, 31 free parameters; competing solutions *K* = 3, BIC = 10,977.02; *K* = 5, BIC = 10,915.10; Figure 6a). AIC favored larger, less parsimonious solutions and did not turn over within the range tested (*K* = 4, AIC = 10,676.49; *K* = 5, AIC = 10,635.66; *K* = 6, AIC = 10,564.20), so we retained the more conservative BIC-selected four-state model. Two states were dominated by broader feature-based looking, which were consistent with a profile of spatial narrowing (subgoal 2). The first, the *size-guided* looking state, was most likely to emit a stop while looking directed at size-similar objects; the second*, the low-similarity* looking state, was characterized by looking at a variety of objects with low similarity to the target in both color and shape. These first two states can be understood as a single subgoal in which the objective is to triangulate the likely spatial location of the target, without requiring identification of the target itself. Two other states were dominated by narrower looking at target-matching objects (subgoal 3): the *conjunction looking* state emitted mostly stops with looking at objects with target-similar size and color, and the *target verification* state was dominated by looking at the target itself. These two states appear to capture a subgoal in which the objective is to identify the target itself.

**Figure 6.**
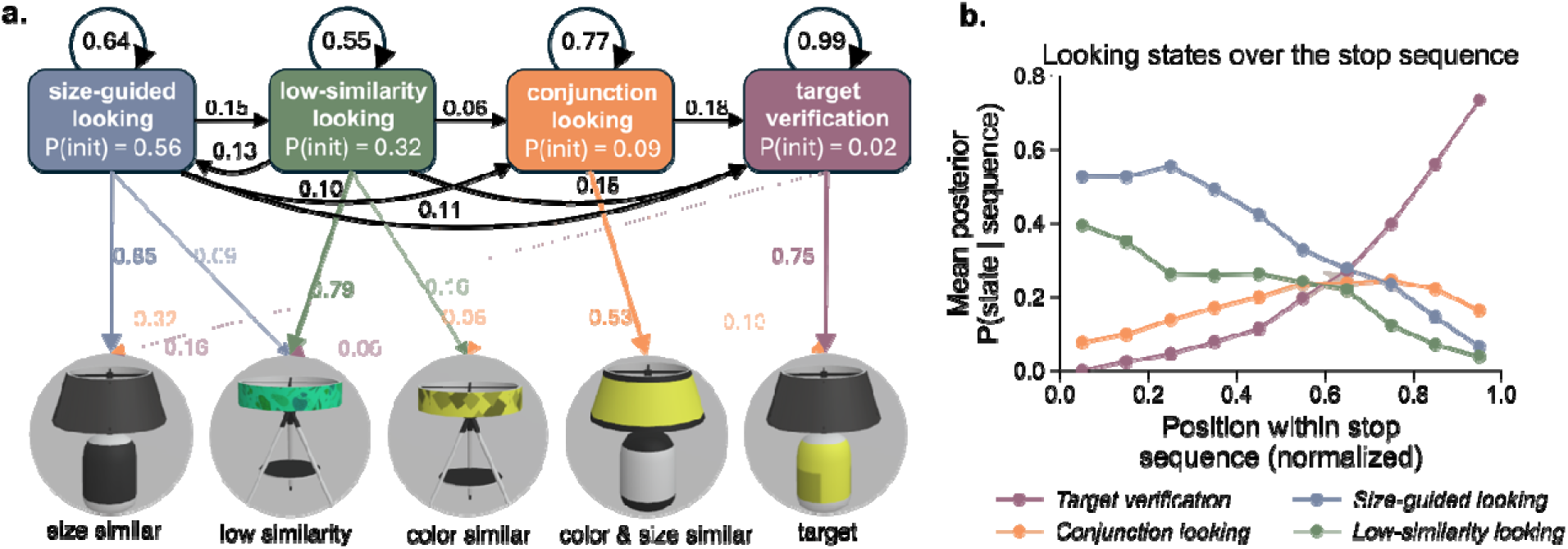
**(a) Best-fitting hidden Markov model.** The model identified four states: two broader feature-based looking states (size-guided looking, low-similarity looking) and two target template-matching states (conjunction looking, target verification). Black arrows show transition probabilities between regimes, and P(init) is the probability of beginning a trial in each state. Colored arrows show, for each state, the probability that a stop emitted in that state was of a given gaze-category type. **(b) Posterior probability of each state across the search sequence.** Mean forward–backward posterior P(state | sequence) as a function of normalized position within the object-stop sequence. Points show the across-participant mean per bin; shaded bands denote 95% confidence intervals.

If the four states characterized the subgoal 2 to subgoal 3 transition over time, as a broad-to-specific search strategy would predict, they should systematically occupy different portions of the trial. The fitted HMM allowed us to test this prediction, as it estimates the likelihood of each state generating each stop; Figure 6b plots these probabilities across normalized within-trial positions. As predicted, trials opened almost exclusively in the two broad search states (initial-state probabilities: size-guided looking = .56, low-similarity looking = .32, vs. conjunction looking = .09 and target verification = .02), and of the two, size-guided looking was reliably more prevalent than low-similarity looking at every decile of the sequence (paired t-tests on logit-transformed trial means, all *p*s < .001, Holm-corrected).

To trace how these opening states transitioned to the narrower ones, we modeled the logit-transformed state probabilities using a regression with states, within-trial position (linear and quadratic terms), and their interactions, and a by-participant random intercept. The four states followed distinct trajectories (omnibus states × position interaction: χ²(6) = 5,546.91, *p* < .001), and a model including quadratic position terms fit reliably better than a linear-only model (χ²(4) = 162.53, *p* < .001; ΔAIC = −154.53, ΔBIC = −123.58). The shape of each trajectory matched the predicted arc from broader to more specific information. Both broad looking states, which dominated the early periods, declined over time (size-guided looking: β_linear_= −6.89, [−7.25, −6.52], *z* = −36.97; β_quadratic_ = −6.50, [−7.78, −5.21], *z* = −9.90, and low-similarity looking: β_linear_= −4.66, [−5.04, −4.27], *z* = −23.68; β_quadratic_ = −3.15, [−4.51, −1.79], *z* = −4.55). The conjunction looking state followed a shallow inverted-U trajectory, rising to a peak in the middle of the trial before declining (β_linear_ = 2.62, [2.18, 3.06], *z* = 11.62; β_quadratic_ = −4.62, [−6.18, −3.07], z = −5.83), consistent with prioritization of the second target feature after the early broader states. Finally, as expected, the target verification state rose at an accelerating rate across the trial (β_linear_ = 10.71, 95% CI [10.36, 11.07], z = 58.83; β_quadratic_ = 3.29, [2.03, 4.54], *z* = 5.13), consistent with identification of the target as the final step in search. Together, the time course of search was well described by a small number of discrete regimes that unfolded in a fixed order.

## Discussion

We sought to test the prediction that self-guided search was best characterized as a hierarchical set of subgoals, nested within an overarching goal to locate and pick up an object, such as the lamp in a store. Although people could have chosen to approach each trial in many different ways, we found a remarkable degree of consistency across individuals. People pursued three subgoals as expected, each of which was governed by its own set of attentional priorities which could be derived empirically from eye gaze. In addition, we identified behavioral markers from head and body movements indicative of planning moments that were common across all subgoals, during which information specific to each subgoal was sought. Together the study provides a novel characterization of self-guided search, a natural behavior that we engage in every day, as a hierarchical plan in which multiple smaller subgoals are pursued sequentially. Our results highlight the importance of considering how navigation, decision-making, visual search, and perception are dynamically integrated during natural behaviors and executed through coordinated body, head, and eye movements.

### Behavioral signatures of goal planning are common to all subgoals

In all subgoal segments, we found a reliable behavioral metric of planning: at periodic intervals, participants stopped for longer and made wider-ranging head movements. We interpret these as pause-and-plan behaviors. These moments are reminiscent of vicarious trial and error (VTE) behaviors in rats, which occur at decision points during maze completion ^9–11,14,32,47,53^. In our study, we found planning moments during all subgoals suggesting that the behavioral signatures associated with planning reflect task-generalizable cognitive moments of deliberation in which the sensory environment is sampled to evaluate options for the next action. Our results show that pause-and-plan behaviors in which individuals stop forward motion and swivel their head are more than route planning; they are behavioral signatures of choice moments common across the subgoals in which attention selects current environmental information to aid decision-making for the next motor actions. Critically, the class of information sampled during pause-and-plan moments differed between the subgoals, as would be expected given the different priorities during self-guided search within each part of the environmental context, as discussed next.

### Subgoals define eye-gaze priorities

We sought to identify the most robust division in search subgoals that could be reliably captured across all people in a fully self-determined search task. During navigation, attention was focused on all path options, which occurred at navigational junctures, suggesting that planning stops were dictated by the path options presented by the local environmental context. Action stops, by contrast, focused more exclusively on the current path. During the two subsequent search subgoals, we found that the distribution of attention also followed a coarse-to-fine progression, first selecting many objects based on their spatial location before homing in on target features. Early in the trial, attention was biased towards low target-similar objects or clusters of objects with one target-similar feature. This result is novel within the visual search literature and suggests that cluster information may act as a spatially constrained informational center of mass that helps to attract or repel attention and subsequent body actions during naturalistic search.

In our study, objects were generally more spatially clustered based on size rather than color in all display rooms. Perhaps because of this, we found that objects with higher size-similarity were more likely to be looked at when they appeared in a tighter cluster.

Clusteredness seemingly accelerated the attentional prioritization of target-similar objects. It is true that in our study, the target could always be found within a target cluster, and therefore looking at size clusters might be biased by that prior knowledge. However, the spatial clusteredness of targets within target display rooms varied significantly (see Methods and Supplementary Figure S7), and the fact that every trial involved a novel target precluded the ability to know the exact configuration of the target’s location. Moreover, there was evidence that dissimilar clusters led to mass rejection of objects, suggesting that the effect of clusteredness on attention was not unique to the cluster in which the target was located, but a more general principle of search. Nevertheless, experiments in which the tightness of clusters for different target features is manipulated will be necessary to dissect the role of clusters in visual search. These data are a first exploration of spatial clusteredness as a modulator of feature-based attentional priority, and they suggest that spatial organization becomes increasingly important for search as spaces become naturalistically larger and more complex.

The use of clusters is important because the utility of massed feature information for selection is likely to transcend any particular task. It has already been argued that the unit of selection is less the object and more a group of objects based on a functional field of view in 2D displays^7,54,55^. This work suggests that the functional field of view extends to naturalistic settings with depth information and may include rapid deduction of the spatial organization of objects.

The common information within a cluster can be understood as a larger mass, or ensemble, of target-relevant information that enhances the likelihood of selection. Clustered information may also be more visible to peripheral vision and therefore offer rapid guidance towards regions of potential high interest ^56–58^ and may even supersede single target-feature based attentional priorities in some situations^59–61^.

We note that three subgoals emerged from our task, but this number is unlikely to capture the range of all natural search tasks. The number of subgoals in other tasks will vary depending on the complexity of the search and the environment. Moreover, the granularity of subgoals may be finer than what we describe, given the natural variability of individual and contextual differences. Our specific findings are likely particular to our environment and task. Search for even the same lamp as in the opening example may proceed very differently if it unfolds in a home, thrift shop, or highly organized international furniture franchise. However, the principles extracted are novel and likely to generalize to other naturalistic, self-guided search tasks. Our results reveal the hierarchical nature of naturalistic search, in which informational sampling and decision-making belong to subgoals that move in a coarse-to-fine progression, from navigational demands to target object selection.

## Materials and Methods

### Participants

Eighty students at the University of California, Davis took part for course credit (53 women, 25 men, 2 nonbinary; median age = 19 years, range = 18–24). The sample identified as Asian (*n* = 42), White (*n* = 28), multiracial (*n* = 4), Black (*n* = 1), and Native Hawaiian or Other Pacific Islander (*n* = 1); 4 participants did not report race. Thirteen participants identified as Hispanic or Latino/a. All participants reported normal or corrected-to-normal vision, with any correction made using contact lenses, and susceptibility to cybersickness was screened in advance using the Visually Induced Motion Sickness Susceptibility Questionnaire ^62^. All study procedures were approved by the Institutional Review Board at the University of California, Davis, and all participants provided informed consent before taking part.

### Apparatus and virtual environment

Stimuli were presented in an HP Omnicept Reverb G2 head-mounted display (4,320 × 2,160 px, 114° field of view) with integrated Tobii eye tracking (120 Hz, accuracy < 1°), calibrated with a nine-point routine before each session. Participants stood within a marked floor boundary in a sound-attenuated room and could turn, crouch, and reach but not walk; the headset cable was suspended from a rod to allow free rotation. They moved through the environment by teleporting, aiming with the controller thumbstick and releasing to move to the indicated location in steps of up to 2 m, and selected objects by aiming a beam and pulling the trigger. The experiment was built in Vizard and run through SteamVR and Windows Mixed Reality.

The environment was a 4,216 m² furniture store with 23 rooms (Figure 2a) with five display rooms, each dedicated to one furniture category (beds, desks, couches, shelves, or lamps), and eighteen living-space rooms arranged as realistic living spaces, connected by hallways. All doorways were open, and some rooms had multiple entrances.

### Procedure

Participants first completed two 4-min exploration trials, freely exploring the store from its southwest corner to become familiar with the layout; no items were collectible during exploration. They then completed ten search trials (Figure 2b). Each trial began with a 6 s preview of that trial’s target object in a blocked-off entryway, after which participants were placed at the southwest corner and asked to search for the target, collecting it by aiming the beam at its price tag from within 1 m. Correct selections ended the trial; incorrect selections were signaled and could be retried until the trial time was up; each trial duration was capped at 4 min.

### Targets and analyzed dataset

The ten targets (two per display room) were the orange and purple couches, orange and yellow lamps, orange and pink desks, green and green-patterned shelves, and yellow and purple beds, presented in two counterbalanced orders across participants (Supplementary Figure S1). Trials in which the target was not found within the time limit were excluded. Each participant contributed a mean of 6.9 analyzed trials (SD = 1.05, median = 7, range 4 to 8), 549 in total, completing 85.8% of their eight non-bed trials. Data were only retained from a person if they completed a minimum of four trials by successfully picking out the target within the allotted time. The two bed targets were excluded by design as filler trials, because the bed display room’s doorway was adjacent to the starting point and required no real search.

### Eye-data preprocessing

A fixation was defined as consecutive gaze within an object’s bounding box lasting at least 100 ms. For the two lamp targets, whose thin parts (such as the stand) were too small to reliably measure fixations on with our system, we increased the fixation bound to include fixations on the wall or platform immediately behind or below the lamp. Additional information based on the earlier-gazed object, head position, and gaze direction was used to assign gaze.

### Segregation of the navigation from search subgoals

Each trial started in the navigation subgoal. This subgoal ended at the first stop at which the participant fixated on a non-structural object inside the target display room and then subsequently entered the display room (Figure 2c). Participants occasionally entered and left the target display room without selecting the target (8 trials total, representing less than 1.25% of trials per participant on average). Edge-case distances and stop counts are reported in the Supplementary Materials.

### K-means for planning and action stop labeling

A distinction we sought to make in our analyses is between planning stops, where participants paused to sample the environment and decide where to go next, and action stops, where they advanced toward a previously chosen region. We identified the two in a data-driven manner from stop duration and head movement. Head movement was summarized as the Total Angular Path, TAP = √(Δyaw² + Δpitch² + Δroll²) summed over successive head samples. Within each trial (excluding the first, in which the head movements were mostly due to participants familiarizing themselves with the environment, and the last stop during which participants figured out how to aim the beam using the joystick), stop duration and TAP were min-max normalized and combined into a single 2D magnitude, √(normalized duration² + normalized TAP²). Normalization was within each trial, given the high variability between trials due to different distances to the target display rooms and changes in navigational efficiency as learning progressed. A k-means model (k = 2) was then fit separately within each phase, classifying higher-magnitude stops as planning stops, and the rest as action stops (Supplementary Figure S2). The choice of k = 2 was supported by cluster-validity analysis of the magnitude distribution in each phase: the silhouette score peaked at k = 2 (0.80 navigation, 0.79 search) and declined monotonically for k = 3 or 4, while a single cluster (k = 1) was rejected by the elbow criterion (the k = 1 to 2 drop accounted for 78% and 85% in navigation and search phase of the total within-cluster sum of squares). Planning stops containing no object fixations (0.2% in navigation, 4.5% in search) were treated as neither type. As a manipulation check, planning stops exceeded action stops on stop duration, head TAP, and the number of paths or objects fixated in both phases (effect-coded mixed models, main effect of stop type p < .001; Supplementary Figure S3). This planning/action distinction is used in every analysis below.

### Navigable paths and path-looking detection

To analyze looks toward paths, we detected the walkable paths available at each stop and classified gaze toward them (Figure 2b). Paths were found by casting horizontal rays outward from the head position at 2° intervals over the full 360°. Only rays longer than 12.5 m were treated as valid, since 12.5 m is the mean distance participants traveled between successive planning stops. A plan formed at a planning stop is only meaningful if it guides movement up to the next planning stop rather than only to the immediate next stop. A path had to stay open across that full stretch to count as a navigable option. Next, the distance to the nearest wall along each ray was recorded, and the peaks in this distance profile within a sliding 40° window (step-size 10°) were taken as the center of a path.

Intervening objects were not used to terminate the ray because they often occluded walkable spaces. The width of the path was determined by the number of contiguous valid rays around the peak. Adjacent or overlapping peaks were merged into a single path. Gaze toward a detected path was classified as the path taken if it lay within 10° of the participant’s next movement and as an alternative path otherwise. See Figure 2b for example path detection and path-looking behavior. This classification scheme contained all the observed stop locations, serving as a validation of the path identification.

### Object similarity and feature clusters

Each target display room was designed so that distractors highly similar in size were spatially clustered, whereas distractors highly similar in color were spatially dispersed, thereby dissociating feature-based and cluster-based guidance in the fixation analysis (Figure 5). Similarity ratings were obtained from a separate 107-participant norming sample^63^, and each object’s spatial clustering was quantified with a nearest-neighbor index (1/NNI). Full norming and clustering procedures, exclusions, and the manipulation check are provided in Supplementary Methods.

### Statistical analysis

The spatial clustering of stop locations was measured as the Shannon entropy of stop counts over a 0.5 m grid and tested against a count-matched, random-sampled null distribution for 1000 iterations.

Trajectory reorientation at each stop was the angle between an incoming vector (from the mean position of the previous two stops to the current stop) and an outgoing vector (from the current stop to the mean position of the next two stops), excluding stops with fewer than two valid neighbors on either side or with step lengths below 0.3 m, and was compared between stop types with Mann–Whitney tests and per-subject Wilcoxon signed-rank tests.

For the path-versus-object analysis, looks were apportioned among four categories (path taken, alternative path, target-relevant object, target-non-relevant object), and the share of fixation time at a stop in each category, arcsine-square-root transformed, was modeled with phase, stop type, and their interaction as fixed effects and target identity as a covariate.

For the cluster analysis, whether each same-category distractor was fixated at a search stop was modeled with a binomial generalized linear mixed model (logit link) with fixed effects of size similarity, color similarity, their interaction, and the spatial clusteredness of size and color (separate size-clusteredness terms for high-similarity (rating ≥ 4) and low-similarity (rating < 4) items), and the interactions between each feature similarity rating and spatial clusteredness; the model also included the number of objects fixated in the stop and target identity as covariates and a by-subject random intercept. For the exploratory analysis of object looking during the navigation phase, whether any object within a display room was fixated (0/1) was modeled with a fixed-effects logistic regression predicting fixation from the object’s size- and color-similarity to the current target.

For the action–planning analysis, the gaze similarity between an action stop and a neighboring planning stop was the complement of the Jensen–Shannon divergence between their four-category gaze profiles, modeled across pairs of each action stop with its nearest preceding and nearest following planning stop, with pair type and the temporal distance to that planning stop as fixed effects and stops containing direct target looks excluded.

Because a state is a distribution rather than a single look or an average over a fixed window, one category of objects can be emitted by more than one state, and two states can share a look category yet differ in when they occur and what they lead to. The temporal structure of search was modeled with a categorical hidden Markov model over the sequence of search stops, each stop coded by its single most frequent gaze category (target, high color-and-size similarity, high color similarity, high size similarity, or low similarity), with both planning and action stops included. The number of latent regimes was selected by BIC over K = 2–5, with categorical emission distributions fit with the hmmlearn Python package^64^ (v0.3.0). Regime dynamics were summarized by the forward–backward posterior probabilities of each regime as a function of normalized within-trial position, and tested using first-versus-last-decile contrasts (paired t-tests on logit-transformed posteriors, Holm-corrected) and within-trial Spearman correlations between posterior probability and position.

### Data and code availability

The analysis-ready data and the analysis code that reproduce all results reported here are publicly available at https://github.com/QQ-Wan/ikea-vr-search. The repository includes the per-stop behavioral tables, the object similarity ratings, and the item-level and model input files, together with the scripts for each analysis and figure. Participant identifiers have been replaced with anonymized codes. Raw head-mounted-display recordings are available from the corresponding authors upon reasonable request.

## Acknowledgments

We would like to thank Makayla Souza-Wiggens, Rishit Das, Catherine Halpern, Zoe Haraeng, Eliana Ertsey, Cailey Tennyson, Nathalie Moriarty, and Morgan Peters for help with data collection and discussions. This work was supported by a James S. McDonnell Foundation grant to JJG and SG.

## Notes

### Competing Interest Statement

The authors have declared no competing interest.

