## Supplementary Materials for "Self-guided search in immersive VR reveals active hierarchical planning"

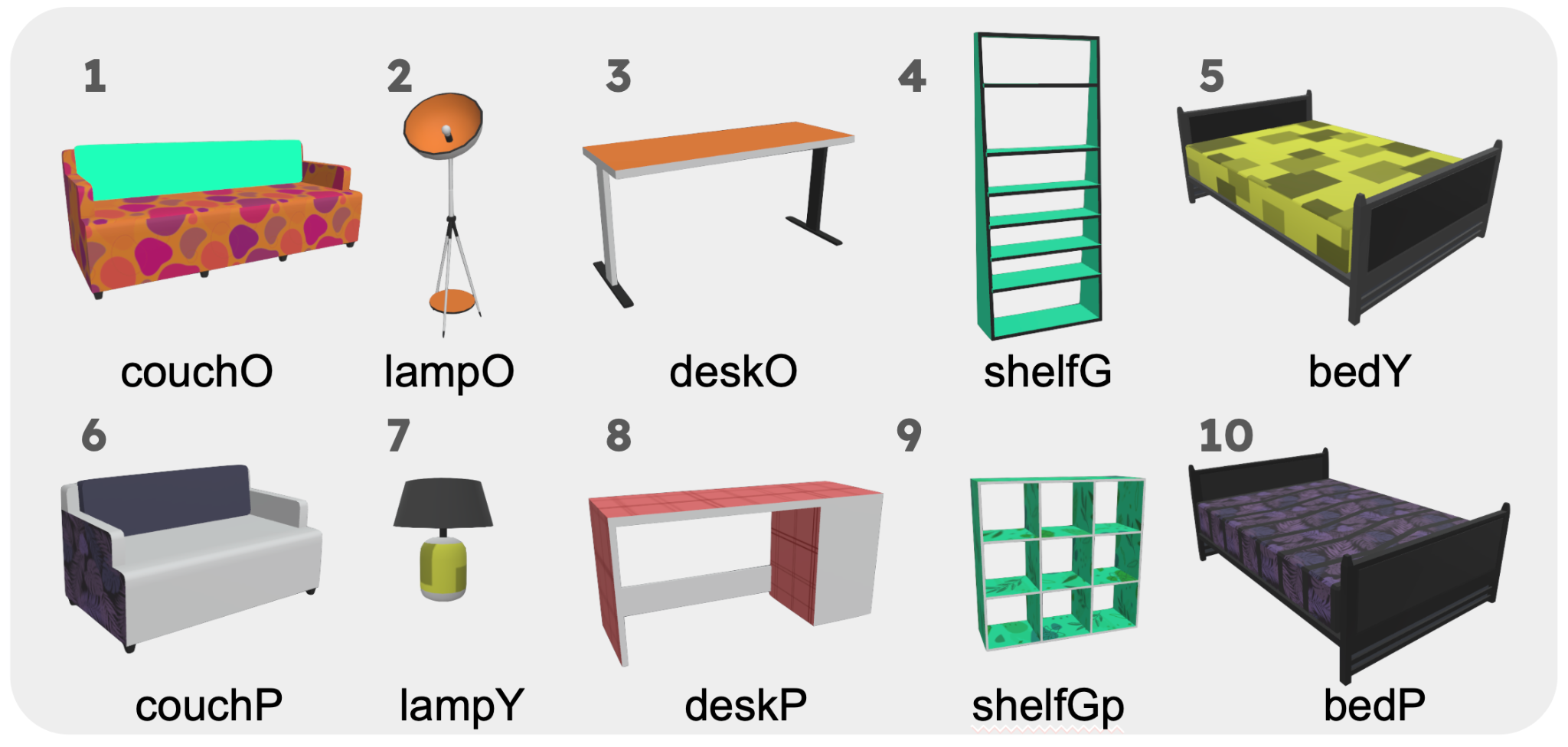

**Figure S1. Targets used in the search trials.** The ten target objects are shown in the order they were presented (the counterbalanced order began with targets 6–10, then 1–5). Each of the five display rooms contained two targets. The bed targets were excluded from all analyses because the bed display room's doorway sat adjacent to the start position and left no navigation demand.

### **Supplementary Methods**

#### **1. Phase-transition edge cases**

On average, 8% of each participant's trials (SD 0.13) contained no stop meeting the phase-transition criterion (a fixation on a non-structural object inside the target display room followed by entry); for these trials the boundary was placed at the first stop inside the display room with a non-structural-object fixation. When the transition stop fell outside the display room, participants were on average 10.13 m (SD 2.08) and 3.95 stops (SD 0.95) from the doorway they subsequently entered; when it fell inside, they were 3.29 m (SD 1.64) and 1.23 stops (SD 1.15) from the doorway they had entered. In the 8 trials (1.46%) in which participants entered and then left the target display room before recognizing it, they spent a mean of 41.08 s (SD 21.74) outside before fixating into the display room again and entering for good.

#### **2. Planning and action stop identification and validation**

Planning and action stops were separated within each phase using k-means clustering (k = 2) on the 2D magnitude, which combines normalized stop duration and head Total Angular Path (Methods). As a check that the resulting labels behaved as intended, planning stops exceeded action stops on stop duration, head Total Angular Path, and the number of paths fixated (Navigation) or unique objects fixated (Search), in both phases. Two effect-coded mixed models (one per phase) with stop type and behavioral-metric type as predictors, the normalized metric value as the outcome, and a by-subject random intercept showed a significant main effect of stop type and a significant stop-type × metric interaction (both *p* < .001).

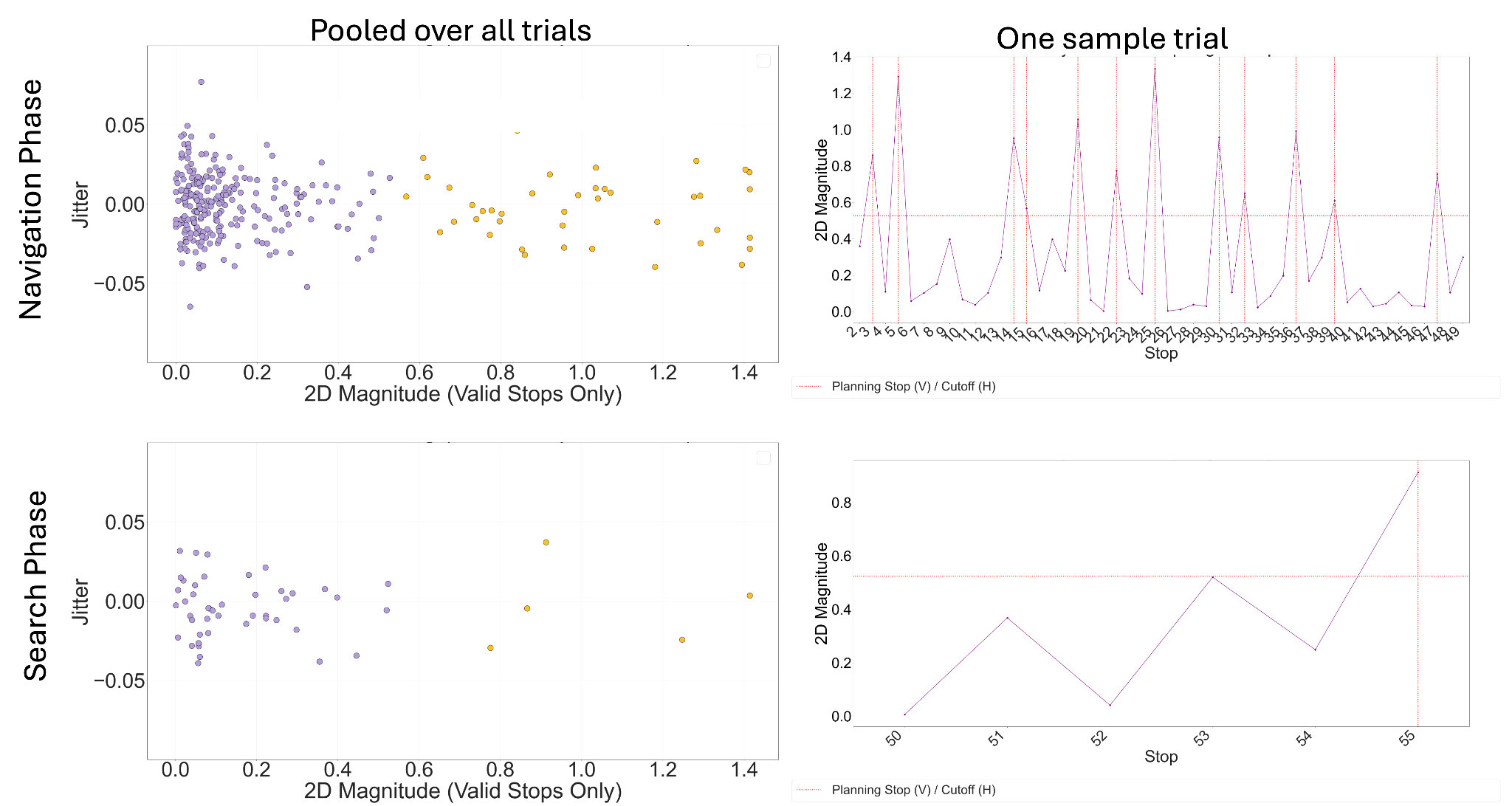

**Figure S2. Identification of planning and action stops.** For one representative participant: k-means segregation of stops by 2D magnitude (left; Navigation above, Search below; stops from all trials pooled, jittered on the y-axis, planning and action stops in different colors), and the 2D-magnitude series across stops within a single trial (right; Navigation above, Search below), with the planning/action cutoff (horizontal line) and the planning stops (vertical lines) marked.

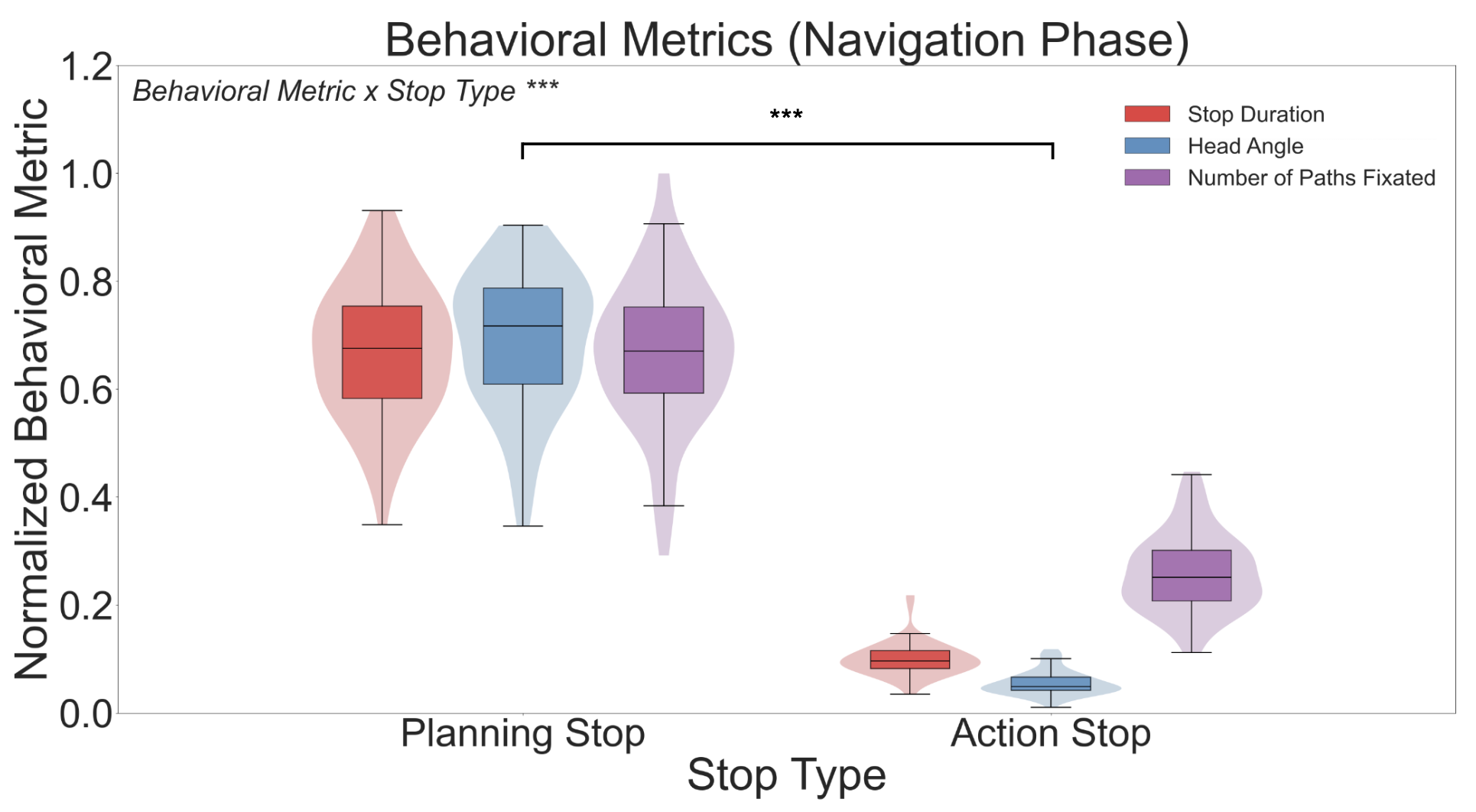

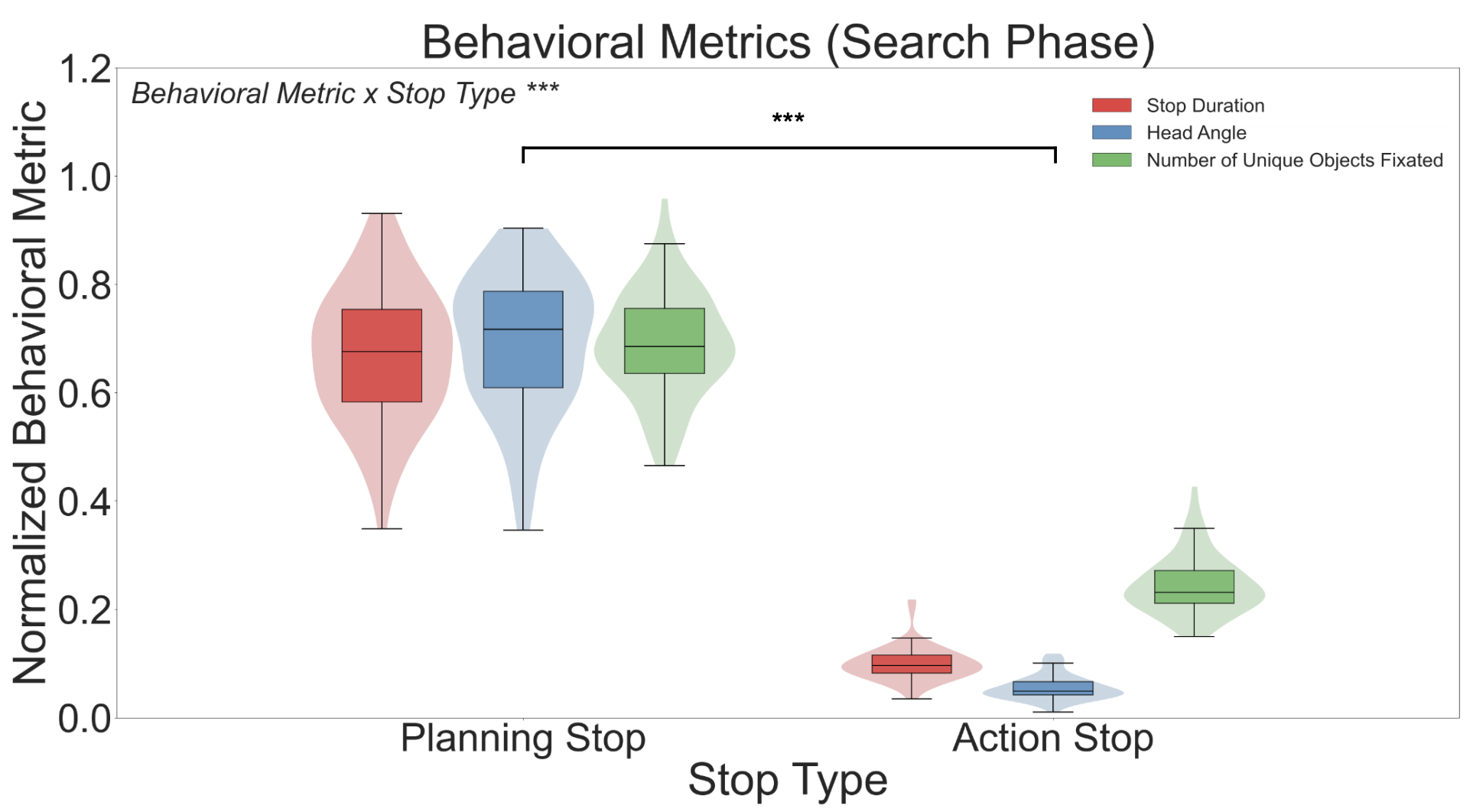

**Figure S3. Manipulation check for the stop typology.** Normalized stop duration, head Total Angular Path, and number of paths fixated (Navigation) or unique objects fixated (Search), by stop type, in each phase (Navigation left, Search right). Planning stops exceeded action stops on every measure (effect-coded mixed models; main effect of stop type and stop-type × metric interaction, both *p* < .001).

#### **3. Object similarity norming and feature clusters**

Similarity ratings were provided by a separate sample of 107 students (online; five participants who failed to indicate maximum similarity for identical objects were excluded). To quantify similarity, this sample rated object pairs (the room's target paired with each other object in that room) on a 6-point similarity scale, separately for shape, size, and color, with at least 21 ratings per pair (Figure S4). The median rating across subjects for each object's similarity to the target was used as the final score. Shape was dropped from all analyses because shape and size ratings were nearly collinear (r(582) = .89) (Figure S5), consistent with the difficulty of differentiating the two when presented together; analyses therefore used size and color. For each target and feature, the high-similarity set comprised all room objects with a median similarity rating of at least 4 (Moderately Similar) on that feature, excluding the target itself; the remaining objects formed the low-similarity set. To quantify each set's spatial clustering, we computed the nearest-neighbor index (NNI): the observed mean nearest-neighbor distance among the set's objects, divided by the mean nearest-neighbor distance expected if the same number of objects were placed uniformly at random over the current object placement area for the display room (200 Monte Carlo samples). We report the reciprocal clustering ratio (1/NNI), which is dimensionless: 1 indicates chance-level spacing and values above 1 indicate objects sit proportionally closer together than expected by chance. The clustering ratio was computed separately for the high- and low-similarity set of each target x feature cell. Each object then inherited, as its spatial moderator in the fixation GLMM, the clustering ratio of the set it belonged to on that feature. As a manipulation check, size-similar objects were more clustered than color-similar objects across display rooms (mean clustering ratio 4.81 vs 2.55; t(9) = 4.57, p = .001; Figure S7).

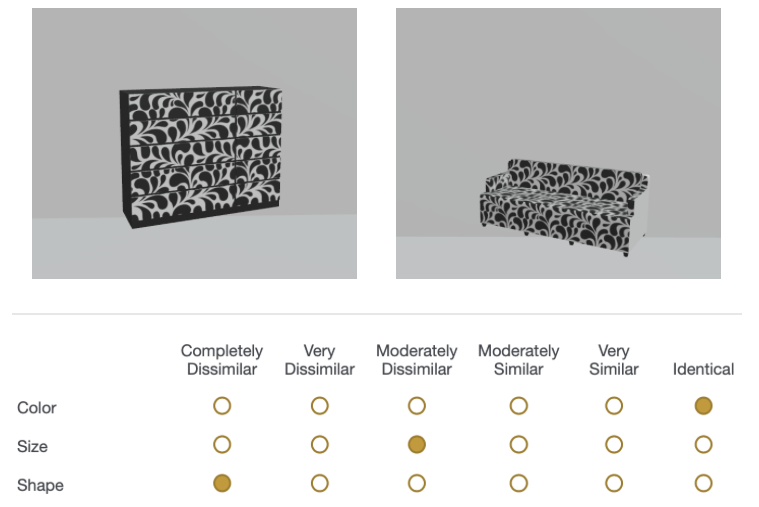

**Figure S4.** *Example trial in the object-target pairwise similarity judgement task*. On each trial, a pair of objects (with one being the target object in the target display room and other object being a non-target object in the target display room) are displayed side by side on the upper side of the screen. On the bottom side of the screen are six-point likert scale options (Completely Dissimilar, Very Dissimilar, Moderately Dissimilar, Moderately Similar, Very Similar, Identical) along with the feature to be judged (Color, Size, Shape) on each row. Participants select only one likert scale option for each feature to indicate their judgement on the similarity between the target object and the non-target object of a certain feature.

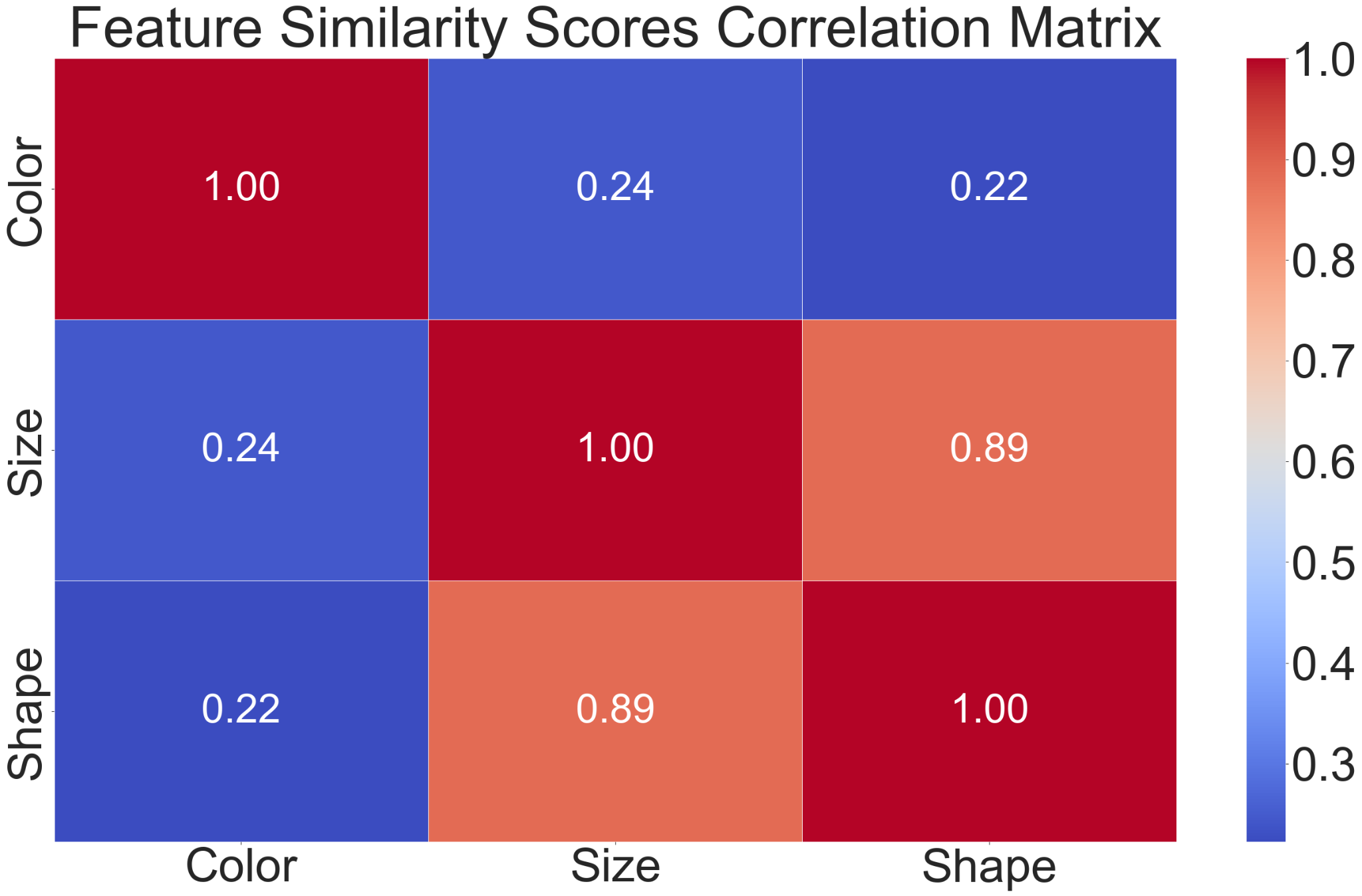

**Figure S5.** *Correlation matrix of object feature similarity scores between features*. The correlation values between each pair of features were computed using the median feature similarity scores obtained from participants’ responses on the object-target pairwise similarity judgement task. Shape and Size features share a high correlation (*r*(582) = 0.89).

### **Supplementary Results**

#### **1. Target features guide eye-gaze during navigation (Subgoal 1)**

Although alternative-path information was prioritized during navigation, we conducted an additional exploratory analysis to test whether the target object was nonetheless attended during this subgoal. Even before participants reached the target display room, the search target already shaped where they looked; as they navigated, participants intermittently fixated objects that resembled the target. We modeled fixation at the level of individual objects, asking whether any object within a display room was fixated (0/1), with fixation predicted from the object's size- and color-similarity to the current target in a fixed-effects logistic regression. Both size-similarity and color-similarity to the current target shaped object looking, with higher similarity predicting higher odds of fixation (size: OR = 6.84, b = 1.92, 95% CI [1.59, 2.26], z = 11.18, p < .001; color: OR = 1.58, b = 0.46, 95% CI [0.09, 0.82], z = 2.46, p = .014). Thus, even in a subgoal where planning priority was wayfinding, the current target's features guided eye-gaze, suggesting the overall goal was maintained across subgoals rather than entirely set aside within each.

#### **2. Robustness of the heading-change measure**

The heading-change analysis in the main text defined the incoming and outgoing trajectory vectors with an averaging window of K = 2 stops. To confirm that the planning-versus-action difference did not depend on this choice, we recomputed the per-participant comparison across a range of window sizes [specify the K values shown]. The effect was unchanged: in both phases, the large majority of participants showed a higher median heading change at planning than at action stops across all window sizes (Figure S6).

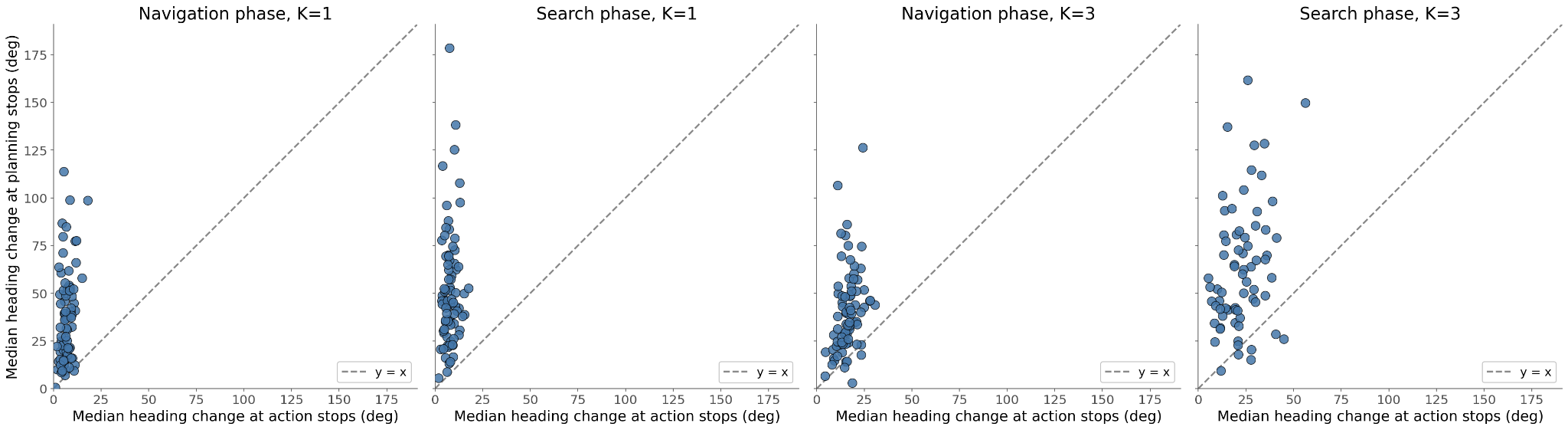

**Figure S6. Robustness of the planning-versus-action heading-change difference to the averaging window.** As in Figure 4, each point is one participant and the dashed line denotes equality, with points above the diagonal indicating a larger median heading change at planning than at action stops. The navigation- and search-phase comparisons are repeated for each averaging window size, K = 1 or 3. The planning-over-action pattern held across all window sizes.

#### **3. Spatial clustering of object features**

Across the ten target display rooms, size-similar objects were more spatially clustered than color-similar objects (mean clustering ratio, 1/NNI, 4.81 for size vs 2.55 for color; medians 4.25 and 2.52; ranges 2.41–7.64 for size and 1.80–3.70 for color). A paired-samples *t*-test confirmed the difference, *t*(9) = 4.57, *p* = .001. Size produced a higher clustering ratio than color in 9 of 10 display rooms; the single exception, Yellow Bed, was a near tie (size 2.41, color 2.51).

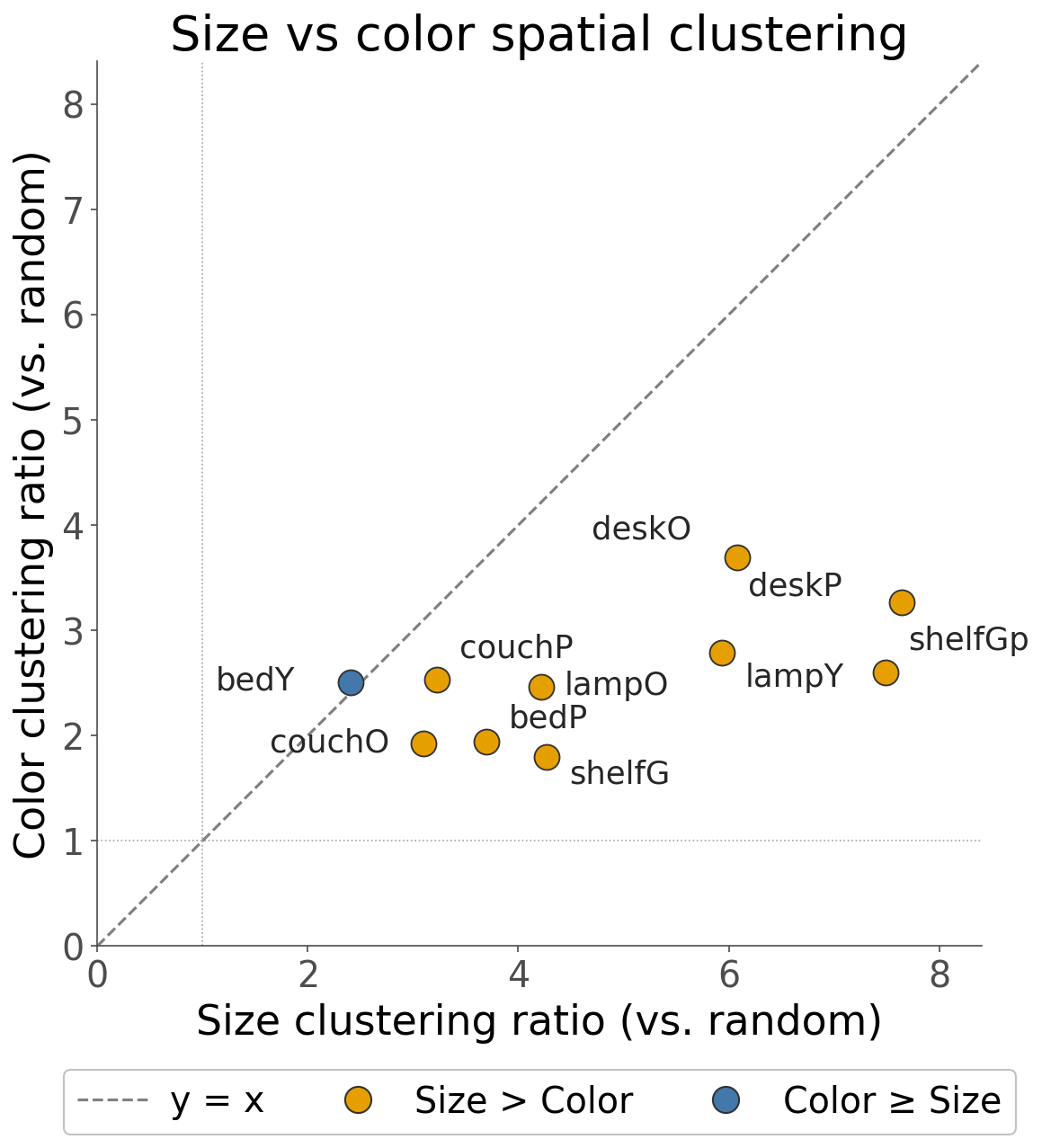

**Figure S7. Spatial clustering of size- versus color-similar objects.** Clustering ratio (1/NNI) for the size-similar and color-similar distractors in each of the ten target display rooms; values above 1 indicate above-chance clustering.

**4. Full model results for fixation inside the target display room (Subgoal 2)**
A generalized linear mixed-effects model (binomial family, logit link) predicted item-level fixation (was_fixated; 1 = item fixated during the stop, 0 = not) from color/size similarity to the target (continuous, centered: size_sim_c, color_sim_c), high- and low-clusteredness indicators for each feature (*_clust_high, *_clust_low), their similarity × clusteredness interactions, a similarity × similarity interaction, the number of items fixated in the stop (centered), and target identity, with a by-subject random intercept.

The model was fit to 56,942 item-observations nested within 1,154 planning stops from 80 participants (overall P(fixated) = .104). The by-subject random intercept variance was small (*σ*² = 0.003, *SD* = 0.055).

We found a significant main effect for feature similarity. Items more similar to the target on size were more likely to be fixated, *OR* = 1.27 (*b* = 0.24, 95% CI [0.22, 0.26]), *z* = 25.15, *p* < .001. The same held for color similarity, *OR* = 1.15 (*b* = 0.14, 95% CI [0.11, 0.17]), *z* = 10.08, *p* < .001. Size and color similarity also interacted positively, *OR* = 1.08 (*b* = 0.08, 95% CI [0.07, 0.09]), *z* = 11.88, *p* < .001.

We found significant main effect for clusteredness. Items in high-size-clusteredness contexts were fixated less, *OR* = 0.86 (*b* = −0.15, 95% CI [−0.22, −0.08]), *z* = −4.18, *p* < .001, and low-size-clusteredness items were fixated less still, *OR* = 0.57 (*b* = −0.56, 95% CI [−0.70, −0.42]), *z* = −8.09, *p* < .001. For color, high clusteredness sharply reduced fixation, *OR* = 0.15 (*b* = −1.90, 95% CI [−2.82, −0.97]), *z* = −4.01, *p* < .001, whereas low color clusteredness did not differ reliably from the reference, *OR* = 0.82 (*b* = −0.19, 95% CI [−0.50, 0.12]), *z* = −1.21, *p* = .226.

We found significant interactions between similarity and clusteredness as reported in the main text. Size similarity's effect was *amplified* under high size clusteredness, *OR* = 1.16 (*b* = 0.15, 95% CI [0.12, 0.18]), *z* = 10.84, *p* < .001, and *attenuated* under low size clusteredness, *OR* = 0.83 (*b* = −0.19, 95% CI [−0.26, −0.11]), *z* = −4.65, *p* < .001. For color, similarity's effect was amplified under high color clusteredness, *OR* = 1.54 (*b* = 0.43, 95% CI [0.08, 0.78]), *z* = 2.40, *p* = .016, but was not reliably moderated under low color clusteredness, *OR* = 0.91 (*b* = −0.09, 95% CI [−0.21, 0.03]), *z* = −1.53, *p* = .126. The color-similarity by color-clusteredness interaction is illustrated in main-text Figure 5c.

As for the covariate in the model, each additional item fixated within the stop increased the odds of fixating a given item, *OR* = 1.15 (*b* = 0.14, 95% CI [0.14, 0.15]), *z* = 54.80, *p* < .001. The intercept of the model is significant too, *OR* = 0.21 (*b* = −1.56, 95% CI [−1.78, −1.34]), *z* = −13.88, *p* < .001.

#### **5. Gaze during action stops anticipates the next planning decision during search**

The main text has shown that participants sampled less information during than the planning stops, but this leaves open the question of what information was sampled at action stops? Because visual sampling inside the target room were largely constraint by the objects category in the target room, it is likely that looking behaviors during action and planning stops share the same information. If true, a question emerges as to whether looking during action stops resembles more to the decision just made - the preceding planning stop or more to the future planning stop, that information was collected and accumulated until the person needs to stop longer to make the decisions.

We tested this with a pairing analysis and found that. Each action stop was paired with the nearest preceding planning stop and the nearest subsequent planning stop. The similarity of the gaze profiles between the action and two planning stops was computed as one minus the Jensen–Shannon divergence (log₂; bounded in [0, 1]) using the distributions of fixation times from the paired stops across the four categories of objects in the display room (objects with high target color-and-size similarity, objects with only high target color-similarity, only target size-similarity, and low-similarity on both dimensions). Stops containing direct looks at the target were excluded since we were interested in looking during the search process. We modeled pair similarity using a linear mixed model that included pair type (preceding vs. subsequent, with preceding as the baseline), the temporal distance to the nearest planning stop (i.e., number of in-between action stops), and a by-subject random intercept.

Action stops resembled the upcoming planning stop more than the just-completed, preceding one (β_following = 0.102, 95% CI [0.061, 0.143], p < .001; Figure S8), indicating a forward-directed bias in gaze. Similarity also decreased with temporal distance to the planning stop, whether that stop preceded or followed the action stop (β = −0.055, 95% CI [−0.066, −0.045], p < .001). Gaze during action stops was thus prospective, anticipating the information that the next planning stop would use rather than lingering on the decision just made. This forward orientation fits the proactive character of looking in natural tasks, where information is sought just-in-time for the next action (see Introduction). It also shows that, within search, action and planning stops sample the same information as a trial progresses, but that new information accumulates until it is only used during the next planning stop to make new decisions. However, because the sampled information is similar across both stop types, the HMM analysis in the main text sets the planning/action distinction aside and treats each stop as a single observation in the search sequence.

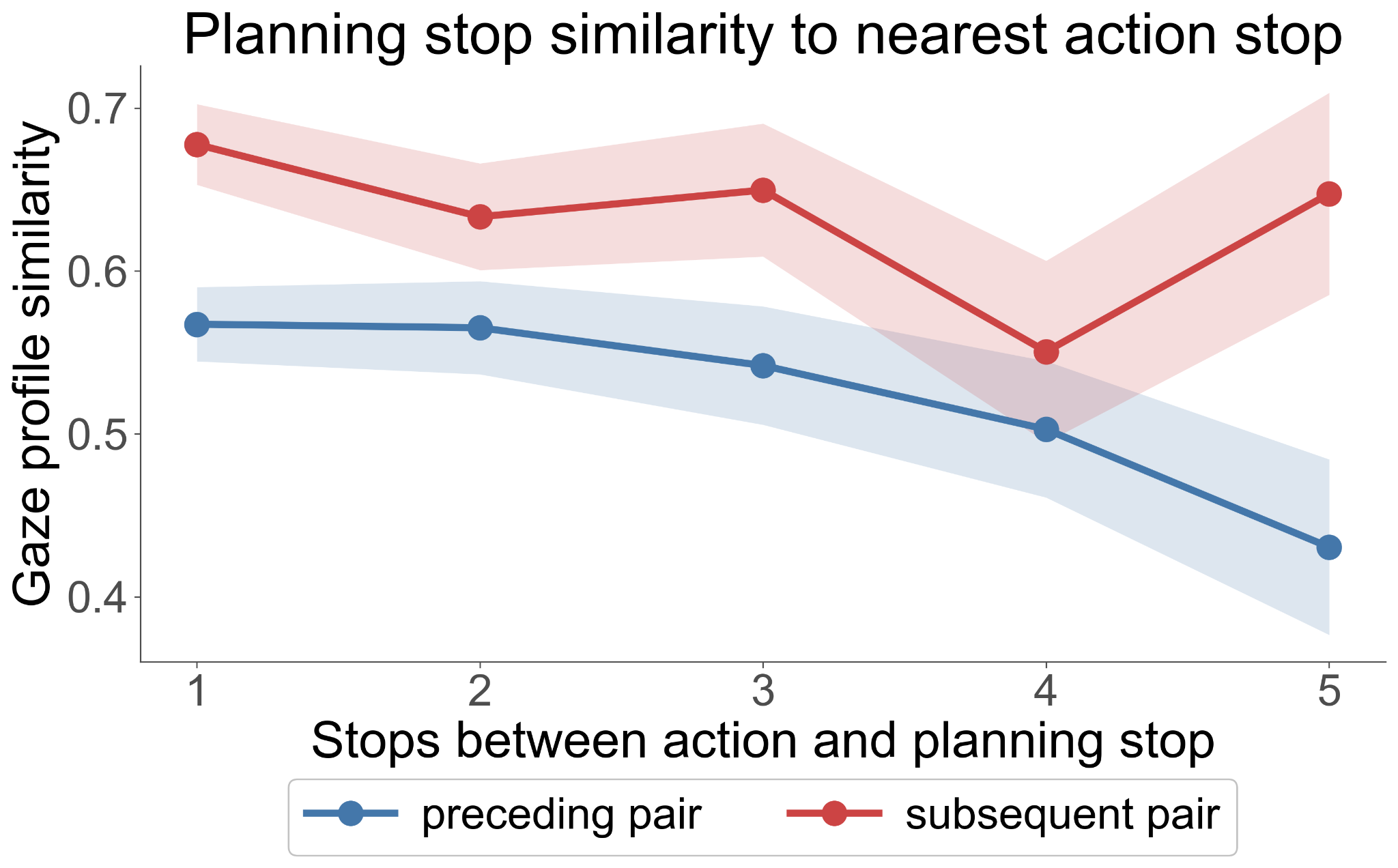

**Figure S8. Gaze-profile similarity between an action stop and its nearest planning stop, as a function of the temporal gap between them.** Similarity is one minus the Jensen–Shannon divergence over the gaze-category profile (bounded in [0, 1]; higher = more similar). Error bars show the standard error of the mean.
